# 3D-Strudel - a novel model-dependent map-feature validation method for high-resolution cryo-EM structures

**DOI:** 10.1101/2021.12.16.472999

**Authors:** Andrei Istrate, Zhe Wang, Garib N Murshudov, Ardan Patwardhan, Gerard J Kleywegt

**Affiliations:** European Molecular Biology Laboratory, European Bioinformatics Institute, Wellcome Genome Campus, Hinxton, Cambridge CB10 1SD, UK; MRC Laboratory of Molecular Biology, Francis Crick Avenue, Cambridge CB2 0QH, UK

## Abstract

Recent technological advances in electron cryo-microscopy (cryo-EM) have led to significant improvements in the resolution of many single-particle reconstructions and a sharp increase in the number of entries released in the Electron Microscopy Data Bank (EMDB) every year, which in turn has opened new possibilities for data mining. Here we present a resolution-dependent library of rotamer-specific amino-acid map motifs mined from entries in the EMDB archive with reported resolution between 2.0 and 4.0Å. We further describe 3D-Strudel, a method for map/model validation based on these libraries. 3D-Strudel calculates linear correlation coefficients between the map values of a map-motif from the library and the experimental map values around a target residue. We also present “Strudel Score”, a plug-in for ChimeraX, as a user-friendly tool for visualisation of 3D-Strudel validation results.

## Introduction

In recent years, technological advances in single-particle electron cryo-microscopy (cryo-EM) have led to an exponential growth of the number of published and archived EM structures^1^. Importantly, since 2018 more than half (54%) of the released Electron Microscopy Data Bank (EMDB)^2,3^ entries have been resolved to better than 4Å resolution. At such resolutions it becomes possible to reliably fit models solved previously, to build *de novo* atomic models and in some cases even to visualise atomic details including the positions of hydrogen atoms.^4,5^

It is widely accepted that validation of structural biology data and models is essential to ensure that the final model is correct and suitable for further use, e.g. to design further experiments or to formulate hypotheses about function, activity, interaction partners, suitable ligands, etc. Methods for validation of atomic models alone are well established as they have been in use in the field of X-ray crystallography for years or even decades^6^. However, another crucial aspect of validation is assessing the compatibility of the atomic model with the experimental data^6,7^ (Kleywegt et al., to be published). This can be done at the level of the entire model and volume or locally to assess smaller-scale features such as individual macromolecules, secondary structure elements, ligands, residues, side chains or even individual atoms. The global fit of map and model is often assessed by calculating the Fourier shell correlation (FSC) curve between the experimental map and a map calculated from the atomic model. Local map-model fit can be evaluated using tools such as atom inclusion^8^, SMOC^9^, EMRinger^10^ and Q-score^11^.

The growing body of high-resolution structures in EMDB and the Protein Data Bank (PDB)^12^ lends itself to systematic structural data mining, which in turn allows for detection of secondary-structure elements in cryo-EM maps^13,14^, extraction of dynamic information from a map alone,^15^ and assessment of protein-protein interfaces^16^.

In this work we have mined the single-particle and helical maps in EMDB determined at resolutions between 2.0 and 4.0Å to produce resolution-dependent, rotamer-specific amino-acid map-motif libraries. We use these libraries in a newly developed method called 3D-Strudel for assessing the fit between a cryo-EM map and these motifs on a per-residue basis. 3D-Strudel calculates the linear correlation coefficient between EM map values for an amino-acid map-motif and the corresponding values in an experimental map around a target residue. The method thus assesses how well the map features around a certain residue resemble those observed in other structures at similar resolution for the modelled residue type, and suggests alternative interpretations of the map where the resemblance is found to be lacking. To facilitate visualisation and interpretation of 3D-Strudel results we have created a plug-in for ChimeraX. We have tested the 3D-Strudel validation method on several map/model pairs including recent SARS-CoV-2 structures and were able to quickly identify model-building errors and regions with generally poor map/model fit.

## Methods

### Overview of the method

The method comprises two separate processes. The generation of the resolution-dependent and rotamer-specific map-motif libraries needs to be carried out only rarely to account for more and newer structures in each of the resolution bands and thus increased frequencies of observation for the rotamers. (We intend to carry out regular library updates at EMDB and to continue to make the libraries publicly available to all users.) The other process is the use of the libraries for validation of a specific map and model, followed by interactive inspection of the results.

### Selection of atomic models and cryo-EM maps

To generate the libraries, single-particle maps (including those with helical symmetry) with fitted atomic models and with reported resolution between 2.0 and 4.0Å were retrieved from EMDB and PDB, respectively. The primary EMDB maps were used for data mining under the assumption that these maps were used to produce the corresponding atomic models.

### Preparation of maps and models for re-refinement

The atomic models were analysed to identify chains with identical sequence and conformation within each entry. The conformational classification was done with hierarchical clustering^17^ as implemented in the scipy Python package using the root-mean-square distance (RMSD) of all non-hydrogen atoms as the metric and using a cut-off of 0.1Å. For each cluster, the chain with the maximum atom inclusion in the map was selected for further analysis. To reduce the resources needed for refinement and map processing, all maps were cropped using a cubic box around the atomic model. The size of the box was chosen such that the distance from any of its edges to the nearest atom was at least 4Å. After cropping, the map origin was set to (0,0,0) and a corresponding shift was applied to the atomic model coordinates, and the fractionalisation matrix was updated accordingly. Refinement dictionaries for any unknown residues were generated using the ACEDRG^18^ and LibCheck programs from the CCP4 suite^19^. There are inconsistencies in the way that fall-off of the power spectrum has been corrected across EMDB entries of the same resolution (B-factor correction). Therefore, we used Refmac5^20^ to normalise this by sharpening or blurring the map until the lowest B-values were between 10 and 15□^2^.

### Re-refinement of atomic models

The atomic models from the PDB were refined using Refmac5. The geometry restraints weight was refined in order to produce a model that has similar bond lengths and angles as the deposited model (RMS differences of their bond lengths no more than 0.02Å and of their bond angles no more than 3°).

The re-refined and original models were compared by calculating the sum of the percent difference in map-model FSC (calculated with Refmac5) and the real-space correlation coefficient (RSCC) (calculated with Phenix^21^). If this number was positive, the re-refined model was selected for further analysis; if it was negative, the original model was used instead. Model quality was evaluated using the Clashscore, Ramachandran outliers, and Ramachandran favourites scores from the Phenix version of MolProbity^21,22^. Entries that pass the validation checks (RSCC > 0.7, Ramachandran outliers < 0.5%, Ramachandran favourites > 90%, Clashscore < 20) were selected for data mining. Detailed model validation statistics for the entries selected for data mining are provided in the **Supplementary Data**.

### Map and model segmentation

For every entry selected for motif mining, separate coordinate and map files were generated for each of its amino-acid residues. For each residue, a cubic box centred at the mean of its atomic coordinates was defined. The box size was chosen such that every atom lies at least 4Å from the nearest edge. To minimise the effect of surrounding atoms, a soft-edge mask based on the distance of map voxels from the model’s sidechain heavy atoms and the C□ atom was applied to the map. The mask is set to “1” if a voxel lies within a distance r_hard_ from any such atom and to “0” if it lies further than a distance r_soft_ + r_hard_ from all these atoms (we used r_soft_=1.0Å and r_hard_=2.0Å). For intermediate distances, the mask value is set according to the formula shown in **Equation (1)**^23^:

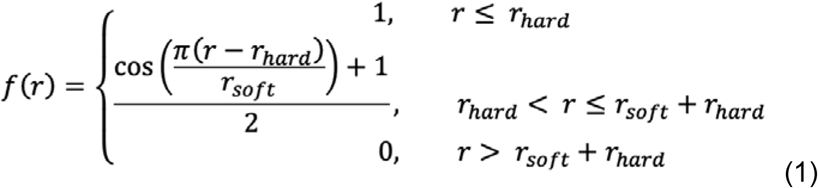

Prior to masking, the maps were oversampled onto a grid with 0.25Å spacing to ensure effective soft-edge masking. The model-map fit for each residue is evaluated using the RSCC (calculated with Phenix) and B-value distribution of all atoms in the model. A residue is considered to have a good model-map fit if its RSCC is greater than 0.7^24^ and the B-values of all its atoms are less than the median plus two times the first quartile of the B-value distribution. Systematically lower RSCC values were observed for negatively charged aspartate and glutamate residues. To ensure the inclusion of a reasonable number of instances of these two residue types we reduced their RSCC threshold to 0.5. The map and model for each residue were saved for further use in the following steps.

### Resolution bands

The segmented maps were assigned the same resolution as the parent map and divided into seven resolution bins: [2.0-2.3>, [2.3-2.5>, [2.5-2.8>, [2.8-3.0>, [3.0- 3.2>, [3.2-3.5>, and [3.5-4.0] Å.

### Rotamer classification

For each residue type, the individual residue models were partitioned into rotamer classes using the peak-occurrence torsion-angle values as defined in the MolProbity penultimate rotamer library^25^. The half-widths at half-height of the torsion-angle distributions were used to define the rotameric intervals.

### Map-fragment superimposition, normalisation and averaging

A reference model consisting of four atoms (C, C□, N, C□) was used to ensure that all segmented maps of a given rotamer were in the same orientation (N-C vector along the X axis, C□-C□ vector along the Y axis). The map fragments were then superimposed in two stages using ChimeraX^26^. First, all the atomic models of the same rotamer were superimposed on the reference atomic model (using their C, C□, N atoms) and the resulting transformation matrices were then applied to the corresponding map fragments. Afterwards, all these maps were averaged to produce an intermediate reference map. Second, all individual map fragments were superimposed onto this intermediate reference map. Subsequently, each map was normalised to have a mean value of zero and a standard deviation of one. Finally, all these maps were averaged again to produce the final average map - this is the map motif for the rotamer under consideration.

### Strudel score and 3D-Strudel validation

We propose the Strudel score as a new metric for cryo-EM map/model validation. It is calculated as the linear correlation coefficient between the map values of a rotamer-specific map-motif from the 3D-Strudel library and the experimental map values around a target residue.

Validation based on the Strudel score has been implemented in an open-source Python software package called *threed_strudel*. The program takes three mandatory inputs: a cryo-EM map, an atomic model, and the path to the 3D-Strudel motif library. The validation protocol consists of the following steps:

1. The map to be validated is split into individual maps, one for each residue, using the model-guided segmentation procedure described earlier.
2. The library motifs of all 154 rotamers of the 20 common amino acids are superimposed in turn on the map around each residue. The atomic model is used to obtain an initial superposition using the C, C□ and N atoms. This superposition is then improved by map-to-map fitting, maximising the RSCC between the two sets of map values.
3. The maximum correlation between the target residue’s local map and the map motifs for all rotamers of the same amino-acid type is reported as the Strudel score for that residue.
4. For every residue, the scores with each of the rotamer motifs for the other 19 amino-acid types are also calculated. The best-matching residue type is defined as that of the rotamer with the highest correlation coefficient. If the best-matching residue type is different from the modelled residue (with a score that is more than 5% better than that of the modelled residue type), the residue is flagged as an outlier.

The program produces output in csv and JSON format. For each residue it contains:

(i) the correlations with the top-scoring motif of each of the 20 amino acid residue types; (ii) transformation matrices which can be used to reproduce the alignment of the motifs with map around the target residue; (iii) a flag for outliers (takes the value one for outliers and zero for non-outliers).

### ChimeraX plug-in

To aid the visualisation and interpretation of 3D-Strudel results we have developed a plug-in for ChimeraX called *Strudel Score*. This plug-in has rich functionality (described in the *Results* section) to facilitate detection of possible model errors and to suggest alternative interpretations. Any model-building issues can be addressed using the ISOLDE plug-in^27^.

## Results

### Calculation of map motifs for all rotamers of standard amino-acid residues

We have mined the EMDB archive to derive map-motif libraries for all 154 rotamers of the 20 common amino acids^25^ in seven resolution bins (**Figure 1**). Libraries for maps with resolution better than 2□ were not derived due to the small number of EMDB entries in this range. An attempt to derive motifs using a wider high-resolution bin (1.5-2.3□) yielded poor results with very high-resolution entries scoring poorly.

**Figure 1.**
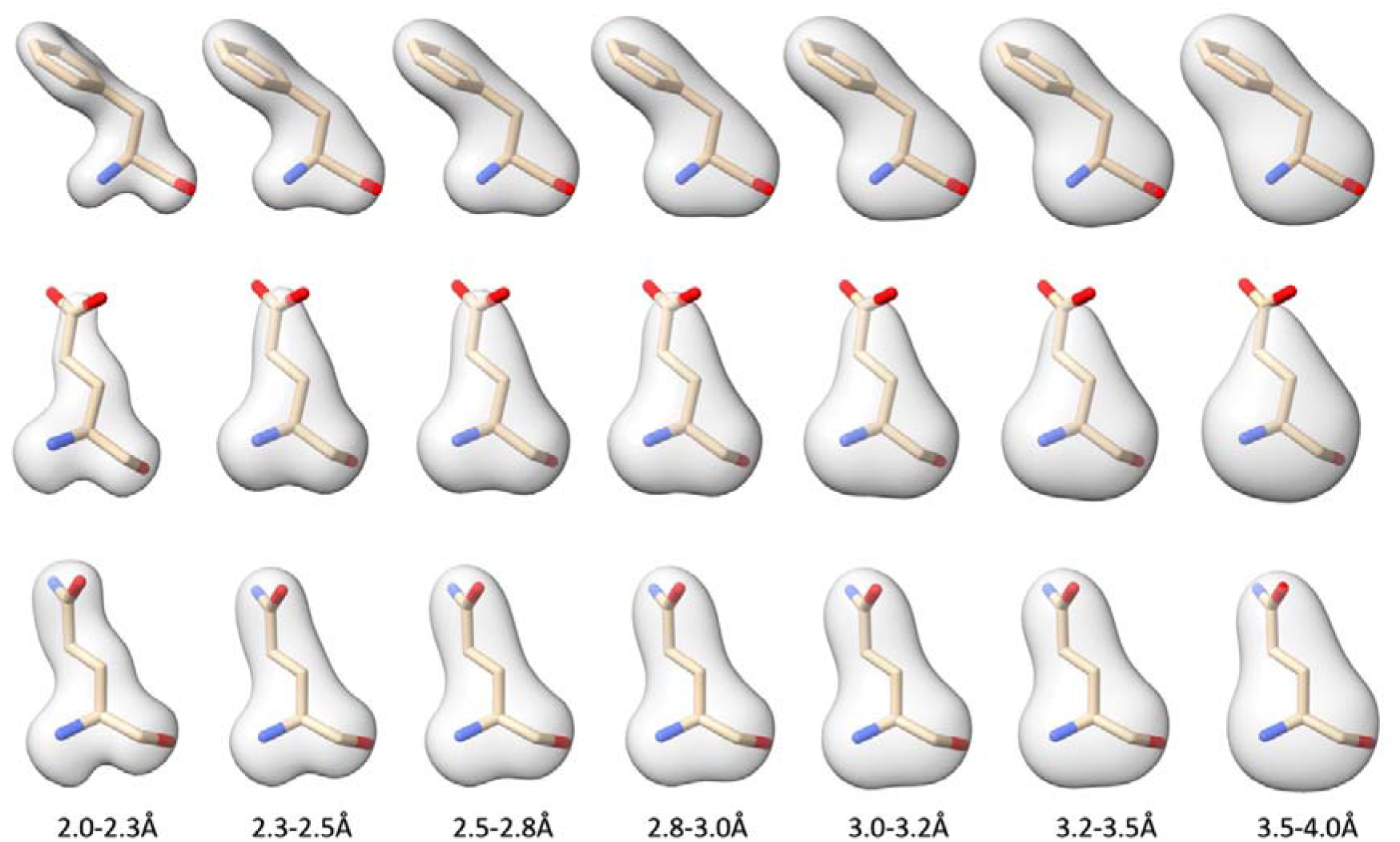
Examples of Strudel map motifs for the seven resolution bands. Top row: motifs for the “m-85°” rotamer of phenylalanine. Middle row: motifs for the “mt-10°” rotamer of glutamic acid. Bottom row: motifs for the “mt-30°” rotamer of glutamine. This clearly shows that the features observed in maps of negatively charged groups differ considerably from those of neutral groups at the same resolution.

This is probably due to their atomistic features which were not well represented in the library motifs as these were largely derived from 2.0-2.3Å maps. We will add bins at higher resolution when more structures in this regime become available and test the applicability of Strudel validation for such structures again at that time.

Deriving the map motifs proceeded in two steps: entry selection and preparation and map-motif generation. In the first step, we selected all maps determined at resolutions between 2.0 and 4.0Å for which fitted or built models were available from the PDB (on October 10, 2021, this search yielded 5239 PDB/EMDB entry pairs). As we intended to use the atomic models to identify map regions around individual residues, a good fit of model and map was essential. However, not all models in the data set have been refined with state-of-the-art approaches and therefore we re-refined the atomic models from the PDB against the primary EMDB map with Refmac5 and assessed the results with MolProbity and Phenix. This resulted in improved map-model FSC and average RSCC for most entries (**Figure 2**). We subsequently selected 3013 map-model pairs that passed our validation checks (see *Methods*) for the motif-generation process.

**Figure 2.**
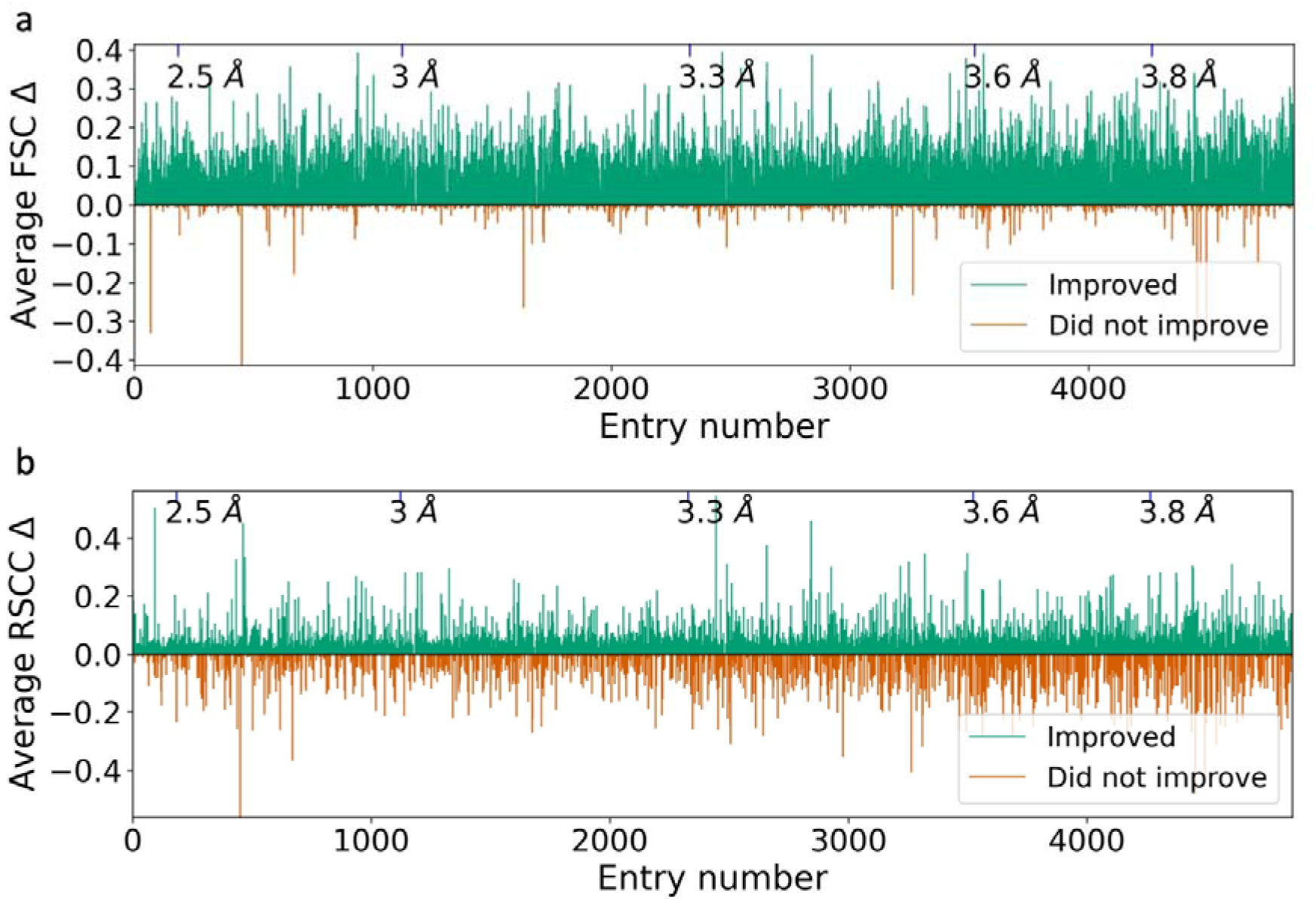
Changes in global map/model Fourier shell correlation **(a)** and real-space correlation **(b)** between cryo-EM models as deposited in the PDB and the same models after re-refinement with Refmac5. Each entry is shown as a teal bar (re-refinement improved the map/model fit) or a red bar (re-refinement made the map/model fit worse), representing the value after re-refinement minus the value for the deposited model. The entries are sorted by resolution. The re-refinement was carried out (1) to reject some models from the 3D-Strudel motif calculations and (2) to improve the fit of model and map for the entries that were selected for the motif generation.

The motif libraries for all resolution bins are openly available for download (CC0 licence). The libraries are versioned and periodically updated; the current version is 4.0 and was released on October 15, 2021. The library for each resolution band is accompanied by files containing detailed information about the data used to derive the motifs, including the EMDB codes and the number of map fragments used to generate each rotamer’s map motif. The total number of map fragments used to derive motifs for a given rotamer ranges from 10 (the minimum required) to ∼140,000. The number depends on the number of available EMDB entries in the given resolution range and the rotamer frequencies in that set of entries. The libraries for all resolution bands between 2.3 and 4.0Å contain motifs for all 154 common rotamers as defined in the penultimate rotamer library^25^. However, the library for the 2.0-2.3Å band contains only 142 motifs. Twelve motifs could not be derived due to a paucity of observations in that band. However, the EMDB archive is growing rapidly and we hope to be able to include the twelve missing motifs in a future library update.

### Library analysis

For practical use, it is important to know how distinguishable the motifs belonging to different amino-acid types are from each other. To investigate this, we calculated the map-map correlation between each rotamer-specific motif and all other motifs (up to 153 of them) in the same resolution band. Then, for each residue type we averaged their correlations with all motifs of the other 19 amino-acid types. Finally, we calculated the quantity one minus the average correlation for every amino-acid type. This value is a measure of how different the motifs of a residue type are compared to the motifs of all other residue types in a given resolution band. Not entirely unexpectedly, the two residues that are most unique are glycine and tryptophan (**Figure 3**), whereas leucine, lysine and glutamic acid are the least distinctive. **Figure 3** also reveals that motif similarity increases significantly as the resolution decreases, which might somewhat limit the potential usefulness of the libraries at lower resolution.

**Figure 3.**
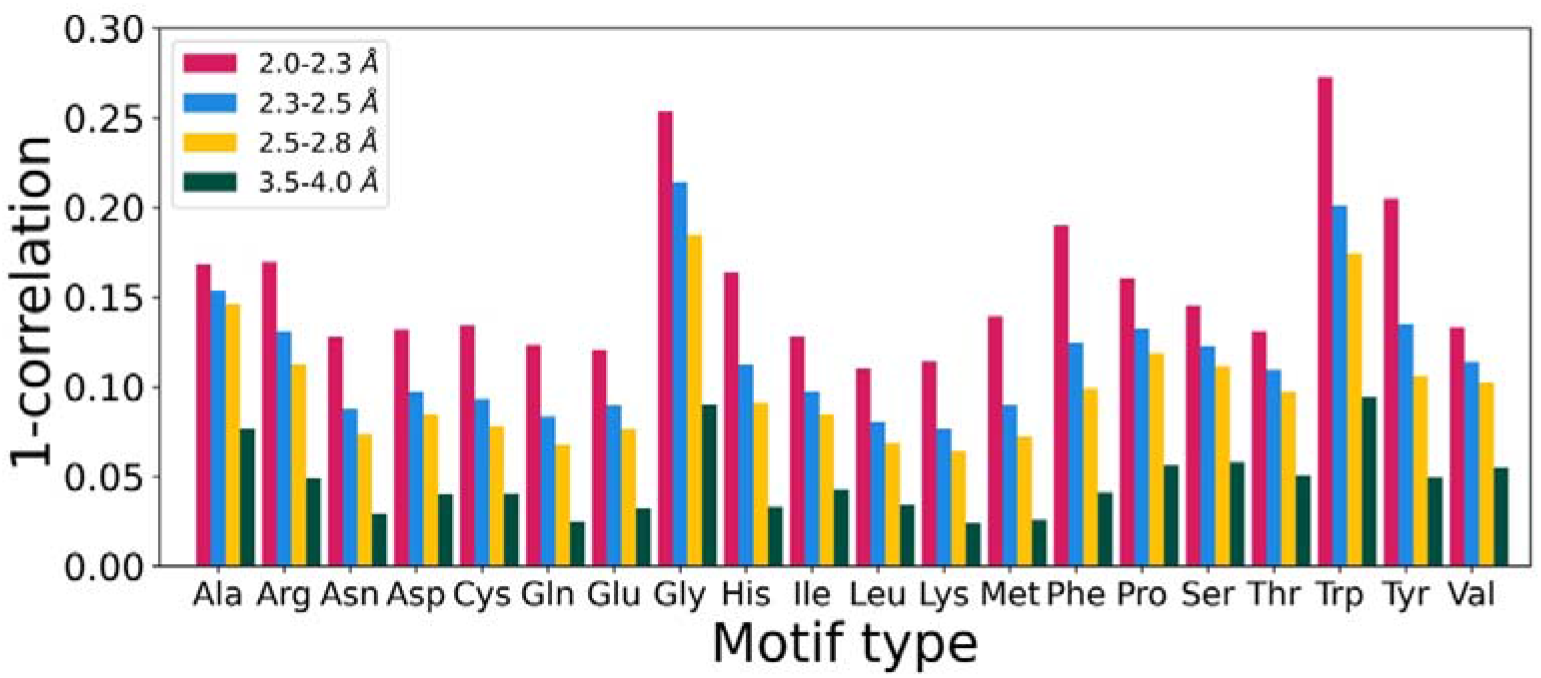
One minus the average correlation between the map values of a residue type’s motifs and those of all other residue types. This quantity indicates how distinguishable the motif maps for a particular amino-acid residue type are from those of all other residue types at a certain resolution.

### Case study 1 - Apoferritin

To demonstrate how 3D-Strudel can be used to detect possible model-building errors, we have applied it to two models (T0102EM054_1 and T0102EM010_1) submitted to the 2019 Model Challenge^28^ for an apoferritin map at 2.3Å resolution (target T0102, EMD-20027^11^). The 3D-Strudel validation results analysis was performed using the Strudel Score plug-in for ChimeraX (program functionality explained in **Figure 4**).

**Figure 4.**
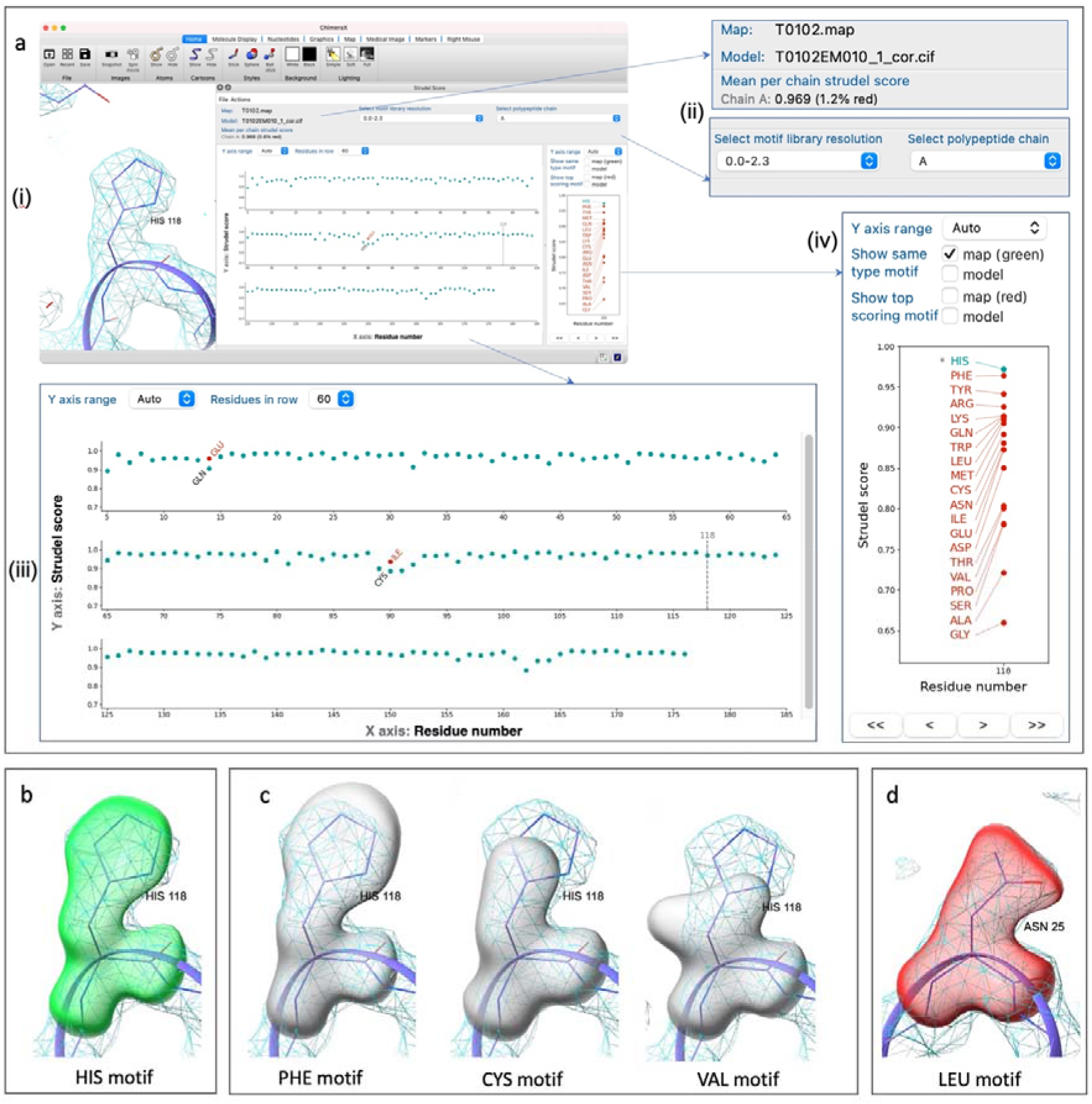
Details of the Strudel Score ChimeraX plug-in. **(a)** Functionality of the plug-in demonstrated using model T0102EM010_1 submitted to the 2019 Model Challenge and the corresponding apoferritin map (EMD-20027) resolved to 2.3 Å resolution. Clockwise from top left: (i) the Strudel Score plug-in integrated in the main ChimeraX window. (ii) General information: map filename, model filename, average Strudel score per chain, percentage of Strudel outliers per chain. (iii) Plots of the Strudel score as a function of residue number. A teal dot represents a residue which scores best (within 5%) with a library motif of the residue type that was modelled there. If this is not the case, the residue is considered an outlier and an additional red dot is shown for the top-scoring residue type. (iv) Upon clicking on a residue in the central graph (residue His A118 in this example) the panel on the right will show the scores for the best-scoring rotamer motif of each residue type. The labels in the right panel can be clicked to show the motifs aligned with the modelled residue and the map: **(b)** in green, the modelled residue type’s motif; **(c)** in grey, the best-fitting motif of any residue type except the modelled one and the top-scoring alternative; **(d)** in red, the top-scoring alternative motif that scores at least 5% better than the modelled residue (if any).

**Figure 5A** shows the Strudel score for residues A65-114 of model T0102EM054_1. Teal dots indicate that a modelled residue is of the same type as the best-scoring motif. If one or more different residue types score significantly (at least 5%) better than the modelled one, the residue is classified as an outlier and an additional red dot is shown for the best-scoring alternative residue type. **Figure 5A** reveals that the region A76-113 of model T0102EM054_1 has many outliers. Visual inspection of map and model reveals that there are two types of outlier. The first type (**Figure 5B, C**) has low scores for both the modelled residue type and for the best-scoring different one, typically less than 0.85. This suggests that the map around the residue is poorly defined or that the residue was built (largely) outside the map. The second type of outlier has high scores, usually greater than 0.9, for one or more residue types that are different from the modelled one (**Figure 5D, E, F, G**). This may indicate that an incorrect residue type was modelled, e.g., due to incorrect sequence information, unexpected mutations, or a register error in the model. For example, the map region where Pro A88 was modelled (**Figure 5D**) scores poorly with the proline library motifs, ranking only 16^th^ out of 20 residue types. Motifs of residue types with long side chains such as Lys, Leu and Met score significantly better. The next residue, Asp A89 (**Figure 5E**), is also flagged as an outlier.

**Figure 5.**
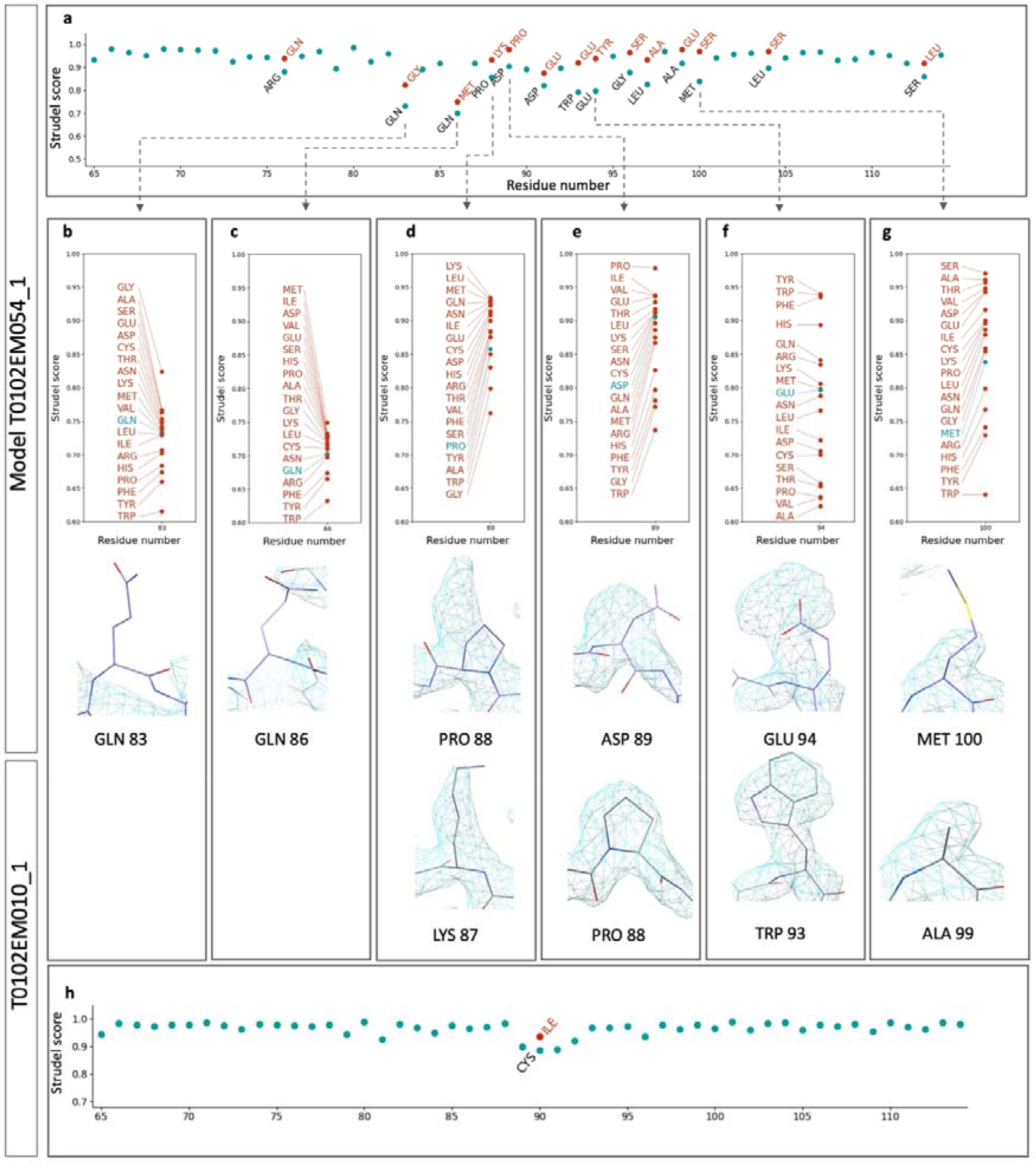
Application of 3D-Strudel for validation. 3D-Strudel results for residues A65-114 of models T0102EM054_1 (**a**) and T0102EM010_1 (**H**) of the 2019 Model Challenge. Strudel-score outliers may arise due to lack of supporting features in the map, for instance, for Gln83 (**b**) and Gln86 (**c**); both the modelled residues and the top-scoring alternatives have relatively low scores (<0.85). Outliers may also be due to incorrectly modelled residue types, in this case a shift by one residue in the assigned sequence in model T0102EM054_1 (panels **d-g**). See text for additional discussion.

Interestingly, here the top-scoring motif is that of a proline residue. Since there is only one proline residue in this region of the sequence (Pro A88) and this was also an outlier, we hypothesised that there could be a one-residue register error in this region. To confirm this, we looked for other outlier residues in this region of the model with easily distinguishable side-chain shapes. An obvious candidate is Trp A93 which scores poorly (**Figure 5A**). The map around residue A94 meanwhile scores best with aromatic residues: Tyr, Trp and Phe (**Figure 5F**) which corroborates the register-error hypothesis.

Model T0102EM010_1 has only one outlier in the A65-114 region. When we inspected the regions discussed above it was clear that this model is correct (**Figure 5H**) and confirms the register error in model T0102EM054_1.

### Case study 2 - SARS-CoV-2 nsp12

An interesting case, which demonstrates the importance of validation and of the ability of Strudel score to detect model-building errors, is EMDB entry EMD-30210 and PDB model 7bv2^29^, one of the first published structures related to SARS-CoV-2. In the originally released atomic model (version 1 of the PDB entry) we detected a register error (A906-920) and a modelling issue (A846-852). Both are located in chain A which corresponds to the RNA-directed RNA polymerase nsp12.

**Figure 6A** shows the Strudel results for the C-terminal region of chain A which has six residues highlighted as outliers. A stretch of outliers usually suggests that the model is poorly fitted, or residue-type mismatches are present. In this case the backbone is fitted well so we checked the top-scoring residue types against the protein sequence which gave a strong indication of a 9-residue register error (**Figure 6B-G)**. When the authors later updated the model (version 2 of the PDB entry), we compared it with our findings which confirmed the register error in the original model. When 3D-Strudel was run with the updated model, all the previous outliers had been resolved (**Figure 6H**). There is a new outlier (His A928), but visual inspection suggests that this is a case of a noisy map rather than a modelling error.

**Figure 6.**
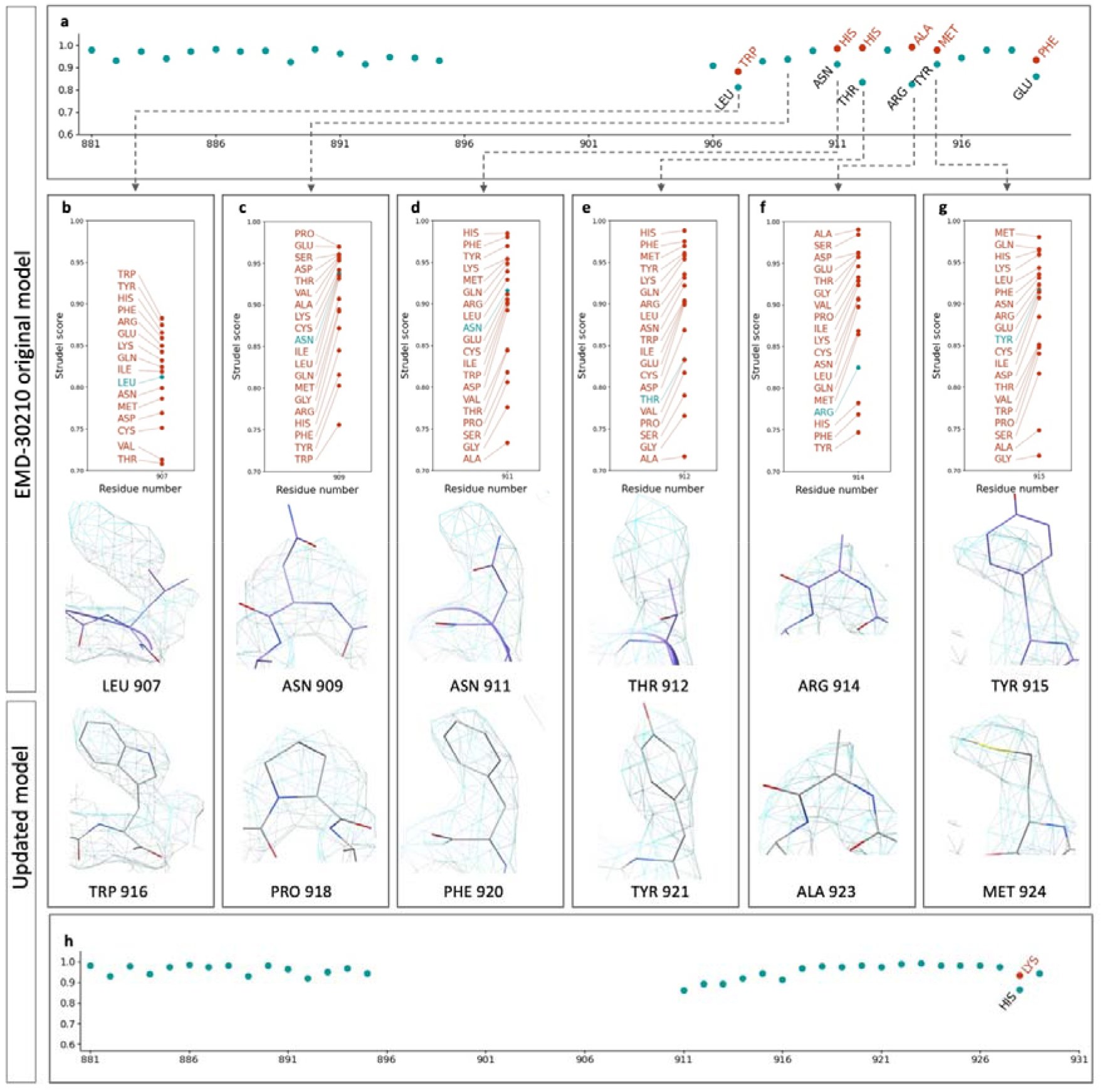
(**a**) 3D-Strudel validation results for the C-terminal region of chain A in the original deposited model of PDB entry 7bv2, built into EMDB entry EMD-30210. (**b-g**) Details of outliers: top row, Strudel scores for each residue type (the modelled residue type in teal, all others in red); middle row, map and model of outlier residues; bottom row, the same location in the map shown with the corrected model. The six outliers suggest a 9-residue register error. **(h)** Strudel scores for the same region as in (a) for the corrected model.

To further investigate the issue, we checked another recently published cryo-EM structure containing nsp12 from SARS-CoV-2^30^ (EMDB entry EMD-30178 and PDB entry 7btf). Strudel validation of the 7btf model (version 2 of the PDB entry) (**Figure S1**) revealed the same 9-residue register error in the nsp12 domain. The presence of such an error in two independently solved structures suggests they might have a common origin and thus we checked the first reported experimental structure of nsp12 from the related virus SARS-CoV^31^ (EMDB entry EMD-0520, PDB entry 6nur). The C-terminal end of the nsp12 protein in the 6nur model (residues A908-920) aligns well with the same sequence in the originally deposited model of 7bv2 (versions 1 and 2 of the PDB entry, which have identical coordinates) (**Figure S2**). This suggests that 6nur may have been used as a starting model for both 7bv2 and 7btf. Strudel validation of EMD-0520 and 6nur shows a stretch of outliers at the C-terminal end of the nsp12 protein consistent with a 10-residue register error (**Figure S3**).

### Strudel results for the EMDB archive

As part of the EMDB Validation Analysis resource^32^ we have calculated Strudel results for all entries between 2.0 and 4.0Å resolution for which a PDB model is available and do so every week for newly released structures. The Strudel results for archived structures can be accessed through the EMDB Validation Analysis resource (http://www.ebi.ac.uk/emdb/va). Results for individual entries can be accessed directly at URLs of the form: https://www.ebi.ac.uk/emdb/va/EMD-30210 (where “EMD-30210” is an example of an EMDB accession).

At the time of writing, Strudel results were available for 5751 structures. The correlation between Strudel scores and resolution, as well as that between Strudel outlier percentage and resolution is fairly low (data not shown; linear correlation coefficients of 0.18 and 0.15, respectively; Spearman rank-correlations of 0.17 and 0.21, respectively). This is to be expected since the Strudel score accounts for resolution by comparing each map against map motifs constructed from structures determined at similar resolution. Nevertheless, the Strudel scores tend to be slightly lower at higher resolution than at lower resolution (median values of 0.956 and 0.971 in the highest and lowest resolution bin, respectively). This could be due to the increased detail observed in individual maps at higher resolution which is averaged out by the motif-generating process and hence not well represented in the map motifs. Strudel outlier percentages, however, are also lower at higher resolution (median values of 2.3% and 6.2% in the highest and lowest resolution bin, respectively). This may reflect the fact that maps at higher resolution are generally of higher quality with fewer poorly defined regions, and errors in sequence assignment are much less likely to occur at higher resolution as well.

As part of our work on the EMDB Validation Analysis resource^32^, and based on community recommendations (Kleywegt et al., to be published) we aim to implement a large number of model-map validation criteria and to carry out a detailed analysis and comparison of their performance in the future. We have carried out only a cursory comparison with the Q-score results^11^ that are also available in the EMDB Validation Analysis resource (data not shown). Strudel scores correlate better with Q-scores than with resolution but the effect is still moderate (linear correlation coefficient of 0.48; Spearman rank-correlation of 0.36; i.e. higher Q-scores generally correspond to better Strudel scores). The correlation is not expected to be very high necessarily as Q-scores are known to be strongly resolution dependent whereas Strudel scores compensate for the effects of resolution through the use of resolution-dependent map motifs. The correlation between Strudel outlier percentages and Q-scores is stronger (linear correlation coefficient of -0.70; Spearman rank-correlation of -0.64), showing that structures with low Q-scores tend to have (many) more outliers than those with high Q-scores. It is clear that the two methods are partly complementary. Whereas the Q-score assesses how well each atom (or, if averaged, each residue) fits the map in the location where it has been positioned, Strudel assesses how well the local map features compare to the average map features observed for the modelled residue type at the given resolution. Since the Strudel score is calculated after map-map superposition (guided by the initial rotamer-map superposition), residues that have been modelled slightly inaccurately may still give good Strudel scores as these do not directly involve the fit of model and map. Thus, we have found entries with both low Q-score values and high Strudel scores which, upon inspection of map and model, turned out to involve models that were slightly rotated from their correct orientation.

## Discussion

The goal of most high-resolution cryo-EM studies is to obtain an atomic model that can be used to provide biological information and insights, and answer questions about function, mechanism, interactions, the impact of mutations, etc. This is a difficult and error-prone task, especially when the structure is solved for the first time and the reconstruction resolution is relatively low. The various examples of register errors (often difficult to identify with other validation methods) that we have detected with Strudel and discussed above illustrate the need for reliable validation tools.

It should be noted that 3D-Strudel uses the atomic model to isolate map regions which have been assigned (possibly erroneously or inaccurately) to individual residues. This limits its ability to suggest alternative interpretations when the backbone is poorly fitted. We recommend visual inspection of the outliers (e.g., with the Strudel Score plug-in in ChimeraX) before considering modifying them to one of the top-scoring alternative residue types. Individual outliers are very unlikely to be due to register errors; only if several outliers occur in a region where the map has good sidechain features need this possibility be investigated.

3D-Strudel validation works best when used with the Strudel motif library of the resolution bin that includes the resolution of the target map. The current set of Strudel libraries has been generated for resolution bins between 2.0 and 4.0Å resolution. We do not recommend using 3D-Strudel for maps with resolution outside of this range. We will add additional resolution bands below 2.0Å when more high-resolution entries become available and the performance of the method has been tested in those resolution regimes.

In this paper we have shown that by comparing the map region where a residue is located with the Strudel motif library it is possible to identify the residue type (assuming there are well-defined features in the experimental map). It has not escaped our attention that in addition to local validation this approach could also be used for protein-sequence assignment/prediction in cryo-EM maps. In a pilot study we were able to successfully predict sequences in maps with resolution between 2.5 and 3.0Å (results not shown).

To date we have derived map motifs for the 20 common amino-acid residues, which are very abundant in the archives and tend to have a few well-defined conformations (rotamers). It may thus not be trivial to apply the same method to other types of molecules that can be encountered in cryo-EM structures (nucleic acids, carbohydrates, ligands).

## Supporting information

Details of structures used to generate the 3D-Strudel map-motif libraries

## Software dependencies and availability

3D-Strudel is a Python-based program that uses several standard Python packages (numpy, scipy, biopython, mrcfile). In addition, it depends on ChimeraX^26^ for alignment of maps and atomic models and for RSCC calculations.

3D-Strudel has been integrated into the CCP-EM package^33^ (https://www.ccpem.ac.uk) and is also available from PyPI (https://pypi.org/project/threed-strudel) and GitHub (https://github.com/emdb-empiar/3dstrudel) under the Apache License Version 2.0.

The Strudel Score plug-in is available from the ChimeraX toolshed (https://cxtoolshed.rbvi.ucsf.edu/apps/chimeraxstrudelscore) and can be installed directly from within the ChimeraX program. The source code of the plug-in is available on GitHub (https://github.com/emdb-empiar/strudel_score). A tutorial is available from the Strudel Score toolshed page as well as from the 3D-Strudel GitHub page.

The 3D-Strudel map-motif libraries are available under a CC0 licence from the EMDB ftp area at https://ftp.ebi.ac.uk/pub/databases/emdb_vault/strudel_libs/.

The 3D-Strudel homepage on the EMDB website (https://www.ebi.ac.uk/emdb/va/strudel) contains more information and will in the future include news about any new releases of both the code and the libraries.

Strudel results for individual EMDB entries can be accessed through the homepage of the EMDB Validation Analysis resource^32^ at https://www.ebi.ac.uk/emdb/va.

## Acknowledgements

We are grateful to our EMDB and PDBe colleagues for useful suggestions and in particular to N. Fonseca for testing the 3D-Strudel software and providing valuable feedback.

We thank A. Joseph and T. Burnley for CCP-EM software support and help with 3D-Strudel integration into CCP-EM. We thank T. Goddard for ChimeraX support and help with Strudel Score plug-in integration into ChimeraX. We are grateful to P. Adams and J. Richardson for help with Phenix and to R. Warshamanage for providing code for map oversampling. We further thank our collaborators in the Wellcome Trust-funded UK EM Validation Network for fruitful discussions: E. Orlova (also for testing the 3D-Strudel software), M. Topf, P. Rosenthal, M. Winn and A. Roseman. We thank C. Lawson and W. Chiu for making the data from the 2019 Model Challenge publicly available and C. Lawson in addition for suggesting the example in case study 1.

AI has been funded by BBSRC (grant BB/P026893/1), the UK EM Validation Network (Wellcome Trust grant 208398/Z/17/Z to P. Rosenthal) and EMBL-EBI. ZW has been funded by the UK EM Validation Network (Wellcome Trust grant 208398/Z/17/Z to P. Rosenthal). Work on EMDB at EMBL-EBI is further supported by the Wellcome Trust (grant 212977/Z/18/Z to AP and GK) and EMBL with funding from its member states. GNM is supported by the Medical Research Council as part of UK Research and Innovation (MC_UP_A025_1012).

