## Supplementary figures and images for "3D-Strudel - a novel model-dependent map-feature validation method for high-resolution cryo-EM structures"

### Supplementary figure S1

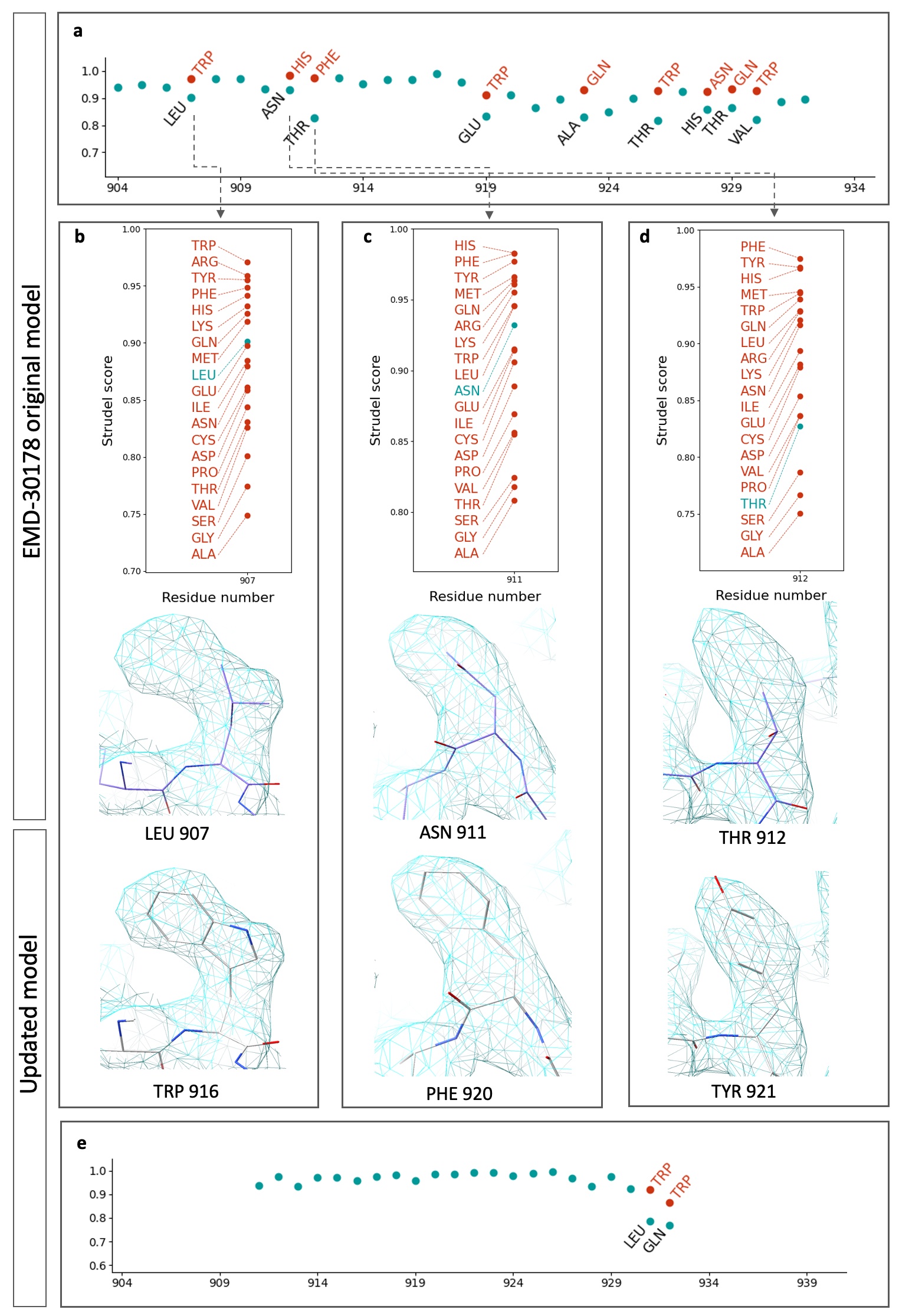

### Supplementary figure S2

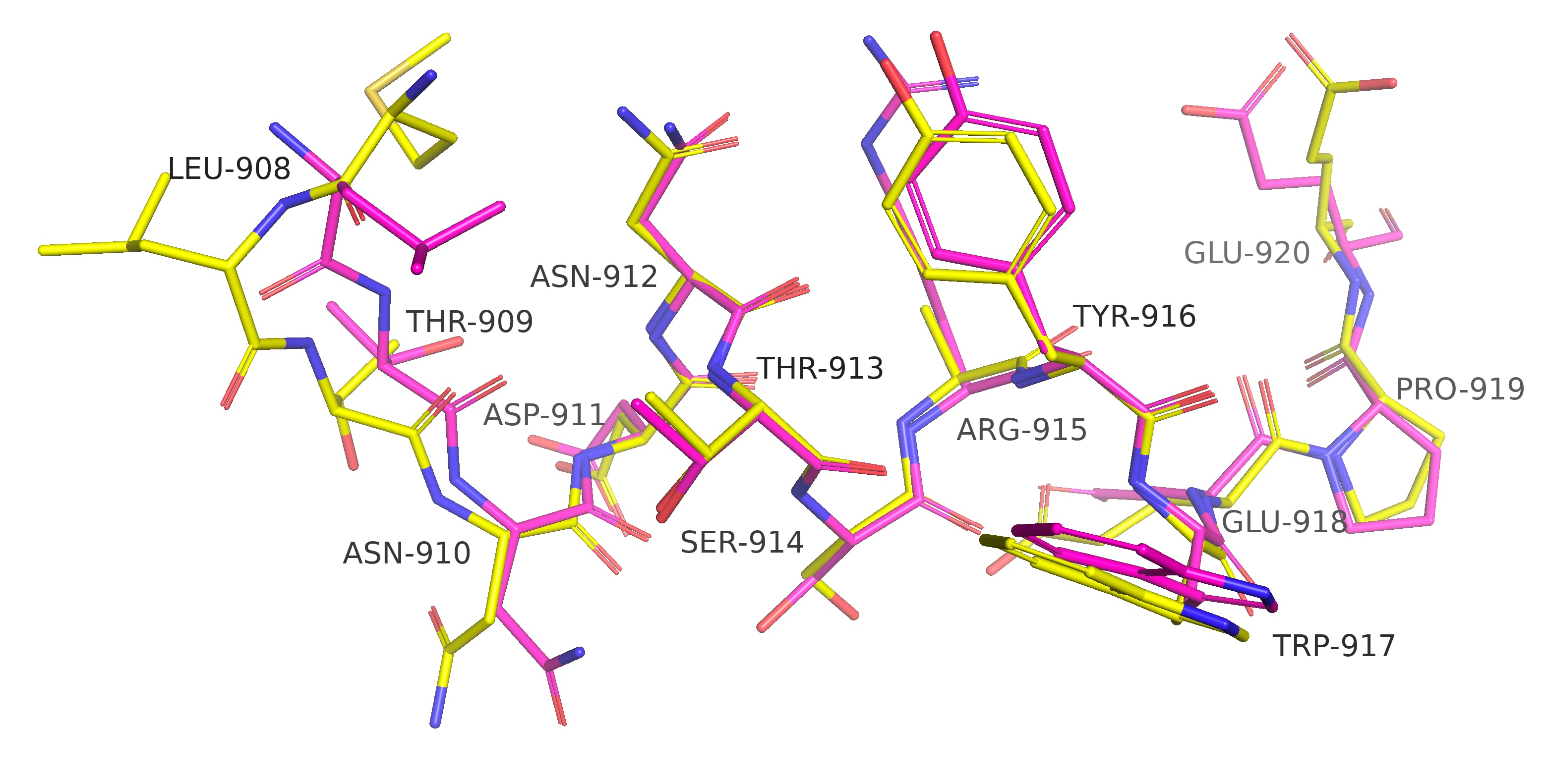

### Supplementary figure S3

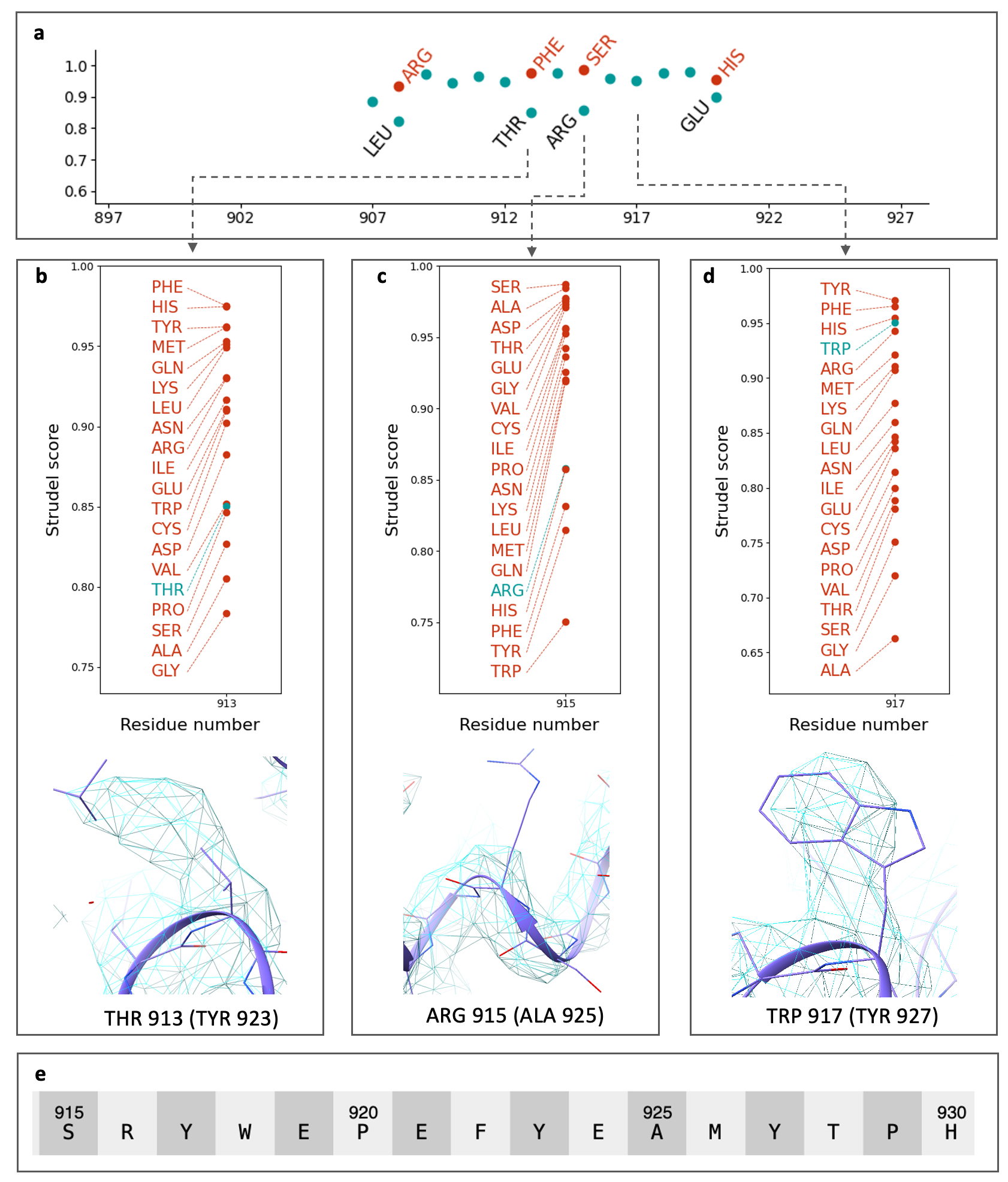
